# Targeting terrestrial vertebrates with eDNA: Trends, perspectives, and considerations for sampling

**DOI:** 10.1101/2024.12.10.627088

**Authors:** Joshua P Newton, Morten E Allentoft, Philip W Bateman, Mieke van der Heyde, Paul Nevill

## Abstract

Terrestrial vertebrates are experiencing worldwide population declines and species extinctions. To effectively conserve remaining populations and species, rapid, cost effective and scalable methods are needed to complement longstanding monitoring methods. Increasingly, environmental DNA (eDNA) based approaches are being used for terrestrial vertebrate biomonitoring within a range of environments. However, as we move eDNA biomonitoring onto land, we are presented with a new set of challenges. This necessitates the development of “best-practice” eDNA sample collection guidelines for terrestrial systems with the purpose of detecting terrestrial vertebrates. To address these needs we conducted a systematic literature review of 143 peer-reviewed papers applying eDNA to terrestrial vertebrate monitoring (excluding Lissamphibia) that were published between 2012 and 2023. We summarize the use of eDNA for terrestrial vertebrate biomonitoring, focusing on study design and field techniques. Over the decade we observe a steady growth in the annual number of publications, with 3 in 2012 and 33 in 2023. The majority of the reviewed studies targeted terrestrial mammals within temperate forest regions. While an equal number of studies focused on a metabarcoding approach to assess community taxon composition and/or species-specific eDNA detection methods, novel uses are increasingly published. These include studies of animal behaviour and population genetics. We record three types of sampling strategies, eight different substrate types and seven different preservation methods, suggesting there is no “one size fits all” eDNA based sampling methodology when detecting terrestrial vertebrates. With a multitude of study aims, across different environments, and target organisms with different ecologies, the standardization of eDNA sampling approaches in terrestrial systems is extremely challenging. We summarize in a table known factors influencing eDNA detection within terrestrial environments. Furthermore, we identify five key considerations to be addressed when sampling for eDNA studies targeting terrestrial vertebrate species, with the aim of guiding decision making.

## Introduction

Terrestrial vertebrates are experiencing alarming declines, with more than 400 species becoming extinct in the last 100 years, and at least 515 on the brink of extinction (fewer than 1000 individuals; Ceballos et al., 2020). Declining populations, extreme losses in geographical range, and localised extinctions are now widespread, even in iconic species such as lions, tigers, and elephants, despite being the focus of significant conservation projects (Alroy, 2015; Ceballos et al., 2017; Ceballos et al., 2020). Biomonitoring is critical for addressing species declines, establishing baseline data to identify and quantify ecological change, and to inform worldwide actions to save wild species (Chiarucci et al., 2011). However, biomonitoring is often hindered by a lack of reliable data surrounding the distribution, habitat association and community dynamics of species (Junker et al., 2020; Rondinini et al., 2011). Long standing biomonitoring techniques targeting terrestrial vertebrates such as direct observations (e.g., field surveys) and traps (e.g., camera trapping, pitfall traps) still play an important role in the collection of this data. However, with ongoing global declines in terrestrial vertebrate species abundance, the development of rapid biomonitoring approaches is now needed to allow for frequent and efficient monitoring.

The number of published studies of terrestrial ecosystems using environmental DNA (eDNA) - DNA derived from environmental samples - has grown rapidly over recent years, because this technology provides a fast and effective method to identify species and hence measure biodiversity (Beng & Corlett, 2020; Nordstrom et al., 2022; Ruppert et al., 2019; Takahashi et al., 2023). Using real time quantitative PCR (qPCR) and droplet digital PCR (ddPCR) it is possible to detect cryptic (Matthias et al., 2021), rare (Koda et al., 2023) or threatened species (Katz et al., 2021), and large-scale eDNA-based biodiversity assessments are possible thanks to the emergence of high throughput DNA sequencing technology combined with DNA metabarcoding (Littlefair et al., 2023). Beyond biodiversity assessments, eDNA can offer insight into many important ecological and biological questions including aspects of animal behaviour (Nichols et al., 2015; Stewart et al., 2018) and plant-animal interactions (Banerjee et al., 2021; Newton et al., 2023). There is also the opportunity to obtain population-level genetic information from eDNA samples (Adams, Knapp, et al., 2019). Furthermore, with vertebrates being the focus of heightened ethical scrutiny and public concern, eDNA offers a non-invasive solution to gathering crucial data on species, without causing harm to animals.

To date, vertebrate eDNA studies have focused primarily on the detection of aquatic species (Ruppert et al., 2019; Sahu et al., 2023; Takahashi et al., 2023) but the application of eDNA-based monitoring for the detection of terrestrial vertebrates is poised to become just as impactful. However, as we move eDNA biomonitoring onto land, we are presented with a new set of challenges due to a vastly different ecology (origin, state, transport, and fate) of eDNA in terrestrial ecosystems. This necessitates the development of “best-practice” eDNA sample collection guidelines that account for the numerous challenges and/or limitations when sampling terrestrial systems with the purpose of detecting terrestrial vertebrates. Here, we conducted a systematic literature review of eDNA studies targeting terrestrial vertebrate species (excluding Lissamphibia) to identify 1) how many studies are published annually, which organisms are routinely being targeted, and what is the geographical and biome distribution of these studies, 2) the applications of terrestrial vertebrate eDNA studies, common sampling methodologies, substrate and sample preservation methods, and 3) best practices in field sampling when targeting terrestrial vertebrates and areas requiring improvement. Based on this, we outline five considerations that need to be addressed when designing the sample collection phase for eDNA studies targeting terrestrial vertebrates, with an ultimate aim to provide a comprehensive review of eDNA sampling approaches and guidance for study design.

## Materials and Methods

In this review, we limit ourselves to the detection of terrestrial amniotes, i.e. all tetrapods excluding the Lissamphibia (e.g., amphibians). We do this as the larval stage of the Lissamphibia and the ecology of the adults blurs the line between terrestrial and aquatic systems; also, the permeable skin of most modern lissamphibians suggests a different rate of eDNA deposition that makes comparison with other tetrapods complicated and worthy of their own review (see Sun et al., 2024). For ease, we hereafter use ‘vertebrate’ to mean ‘amniotic tetrapod’ unless otherwise indicated. In line with Thomsen and Willerslev (2015), we define eDNA as genetic material obtained directly from environmental samples (soil, sediment, water, etc.), without any obvious signs of biological source material. However, we extend this definition to encompass DNA obtained from artificial materials that similarly lack obvious signs of biological source material (e.g., assessing bite marks from clay models; Shaw et al., 2023). They, and we, also excluded studies of other complex samples such as invertebrate derived DNA and faecal DNA studies.

Literature searches were conducted in Scopus and Web of Science in January 2023 for peer-reviewed papers published between 2012 – 2023, using the search terms “Metabarcod*” OR “Environmental DNA” OR eDNA AND in combination with terms specific to the three terrestrial amniote taxa (Aves, Reptilia and Mammalia; Appendix S1). The literature search identified 2,682 unique and potentially relevant publications.

After excluding faecal and iDNA studies, those solely focusing on Lissamphibia species, reviews, ancient DNA studies, government documents, and entirely industrial documents 143 papers remained. For each of the remaining publications, we recorded the following information: study taxa, study country and terrestrial biomes (simplified from Olson et al., 2001), study aims, sampling strategy, substrate type collected, contamination reduction strategies, types of negative controls, preservation methods, target gene region and length (bp) and information of permissions or ethics approvals gained (Appendix S2, Table S3).

Analyses and data visualization were undertaken in R (v.4.0.0; R Core Team, 2024). When multiple methods were used in a study, each method was counted independently for the summary statistics and analyses. To show overall trends in eDNA studies for the detection of vertebrates, an Alluvial plot was created using RawGraphs (Mauri et al., 2017) and spatial illustrations were made in RStudio using Natural Earth data.

### Overview of data

After curation we identified a total of 143 eDNA-based studies focused on the detection of terrestrial vertebrates (but not restricted to terrestrial substrates e.g., the detection of terrestrial mammals from water sources), showing a strong growth in the number of publications since 2012 (Figure 1b). The number of publications peaked in 2023 (33 studies), followed by the year 2022 (25 studies) and 2021 (24 studies), reflecting a rapid increase in outputs of studies over the past decade. The majority of the 143 reviewed studies were targeting and detecting mammals (95 studies - 66 %), followed by reptiles (52 studies – 36 %) and birds (47 studies – 33 %; note, because some studies focused on more than one taxon, the total number of published studies and studies by taxa do not correspond). This taxon bias towards mammals was similar for all continents; however, Oceania produced a roughly equivalent number of bird and mammal studies, while both N. America and Asia produced a roughly equivalent number of reptile and mammal studies. The highest number of studies (60%; Figure 1a) were conducted in North America (55 studies – 38 %), and Europe (31 studies – 22%). Very few studies have been conducted in Africa (6 studies – 4 %) as a whole. Of the world’s terrestrial biomes (simplified from Olson et al., 2001), studies conducted within temperate forest regions have dominated the number of publications (64 studies – 45 %) followed by tropical forest regions (30 studies – 21%). All other biomes are each represented in less than 9 published studies (Figure 1c).

**Figure 1.**
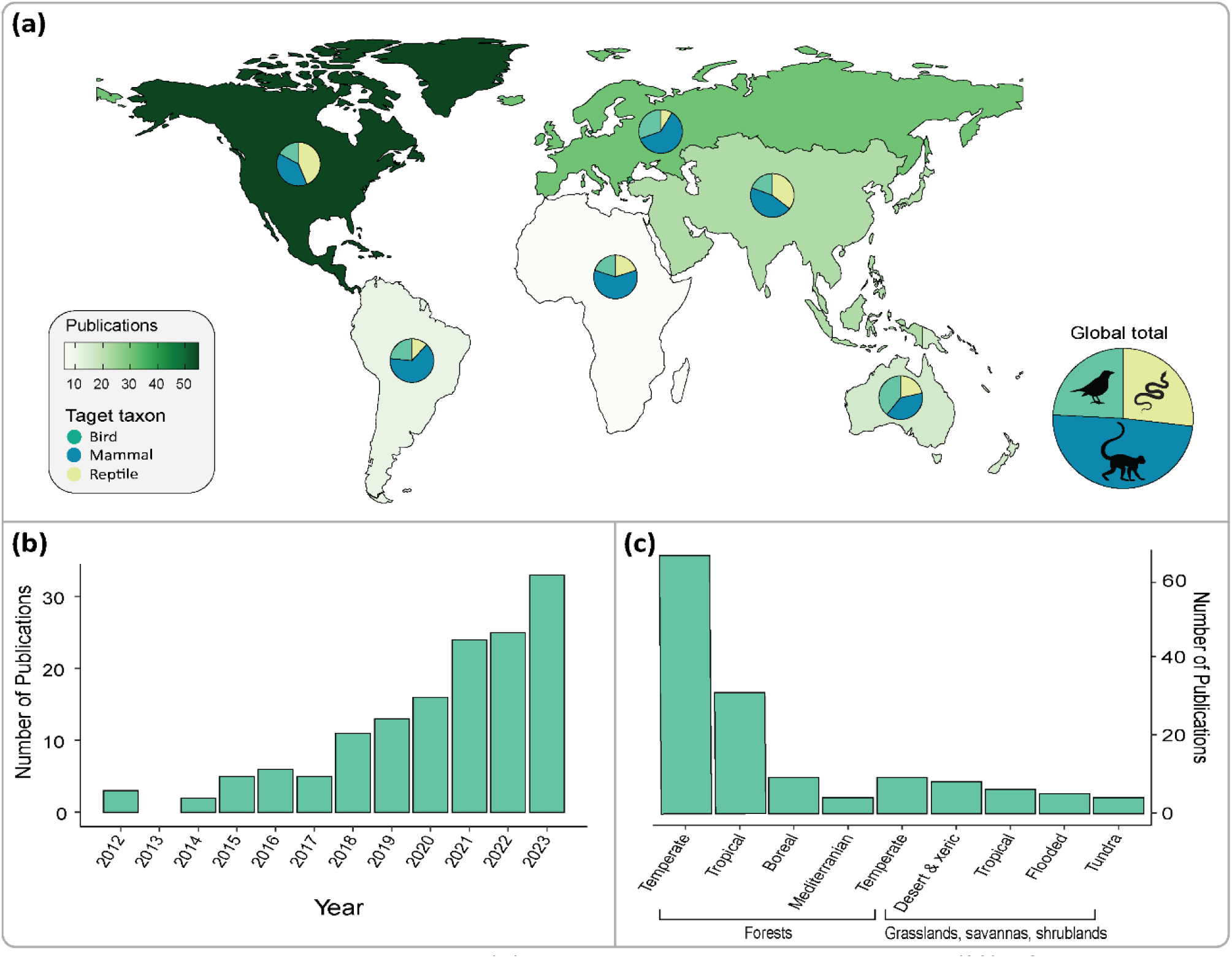
Literature search results: (a) Map illustrating the proportions (%) of studies targeting vertebrate eDNA (excluding Lissamphibia) in terrestrial systems across six continents – Africa, Asia, Europe, North America, South America, and Oceania. The target taxon (Bird, Mammal and Reptile) per region and global total are indicated in the pie graphs. The studies are assigned to country and continent as by where the fieldwork was conducted, as specified in the reviewed papers; (b) Terrestrial vertebrate eDNA publications per year from 2012 to 2023 (excluding Lissamphibia); (c) Number of terrestrial vertebrate eDNA publication (excluding Lissamphibia) for the worlds terrestrial biomes simplified from Olson et al. (2001).

#### Current applications for terrestrial vertebrates

The literature search highlights a wide range of potential applications of eDNA, encompassing biodiversity assessments at various spatial scales (e.g., Broadhurst et al., 2021) and across different land-use (e.g., Ionescu et al., 2022), monitoring vertebrate activity patterns (e.g., M. J. Farrell et al., 2022), the detection or rare (e.g., Priestley et al., 2021), cryptic (e.g., Matthias et al., 2021) or endangered species (e.g., Walker et al., 2022) for better conservation outcomes, estimating species distributions including those of invasive species (e.g., Piaggio et al., 2014; Williams et al., 2018), studies of animal behaviour (e.g., Shaw et al., 2023), and studies of plant-animal (e.g., Newton et al., 2023; Walker et al., 2022), or predator-prey interactions (e.g., Rößler et al., 2020). An equal number of studies have focused on multispecies metabarcoding to assess community composition (61 studies – 43 %; e.g., Figure 2a) and species-specific eDNA detection methods (61 studies – 43 %; e.g., Figure 2b), such as quantitative PCR (qPCR). The remaining, mainly recent, studies go beyond taxon presence- absence data. For example, recently plant-animal interactions were investigated (e.g., Figure 2c), and the use of eDNA approaches to obtain population genetics data (e.g., Figure 2d). These studies, while few (Figure 3a), indicate the maturation of eDNA as a tool in terrestrial ecosystems as researchers begin to utilize this technology to answer more complex ecological questions.

**Figure 2.**
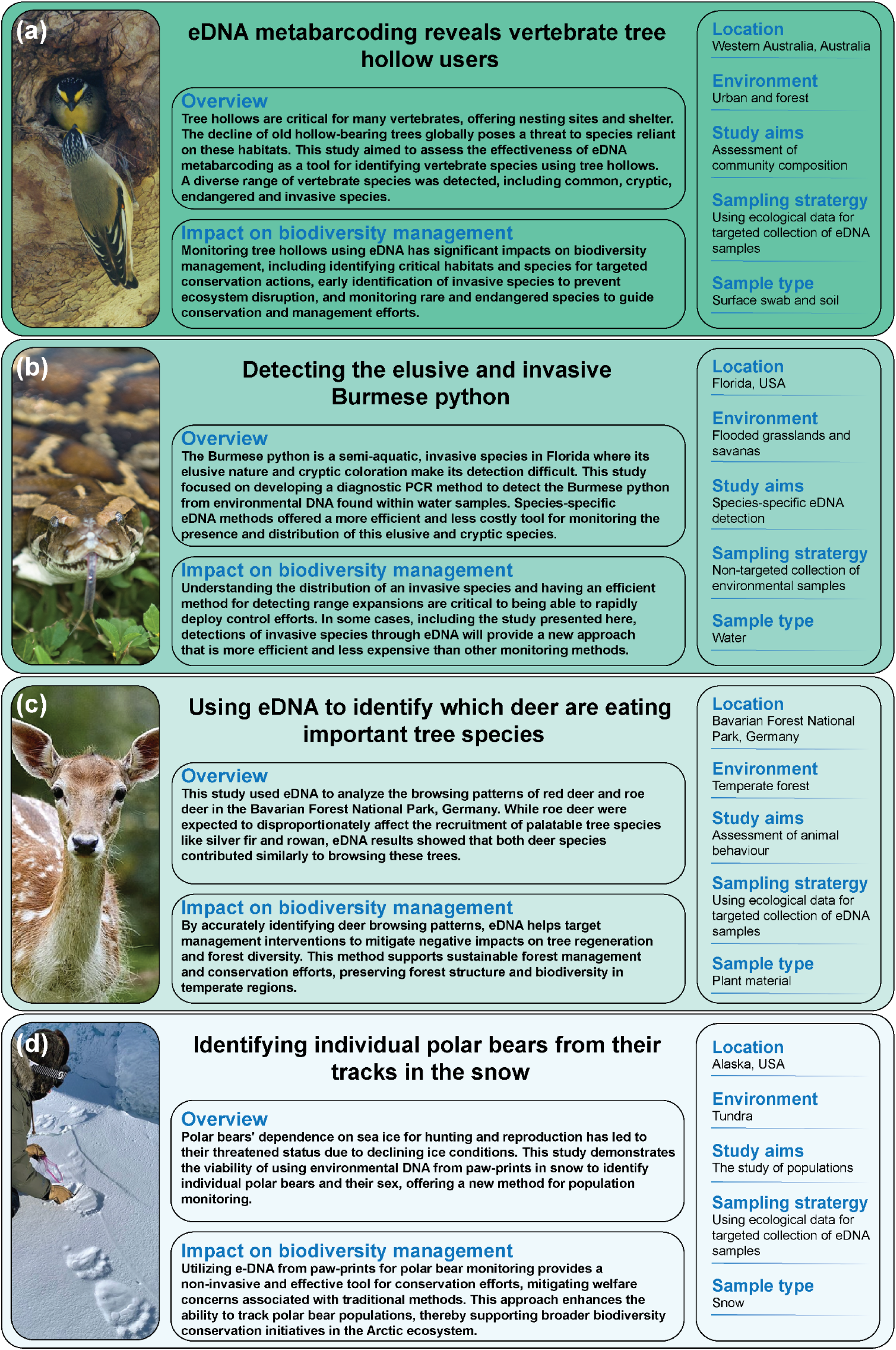
Short case studies highlighting potential uses of eDNA for the monitoring of terrestrial vertebrate species and the impacts on biodiversity monitoring. Cases studies drawn from (a) Newton et al. (2022), <u>photo</u> by G. Bowland (<u>CC BY-SA 3.0</u>), via Wikimedia commons, (b) Piaggio et al. (2014), <u>photo</u> by South Florida Water Management District (<u>CC</u> <u>BY-ND 2.0</u>), via flickr, (c) van Beeck Calkoen et al. (2019), <u>photo</u> by A. Öztas (<u>CC BY-SA</u> <u>4.0</u>), via Wikimedia commons, (d) Von Duyke et al. (2023), photo from Von Duyke et al. (2023) (<u>CC BY 4.0</u>).

**Figure 3.**
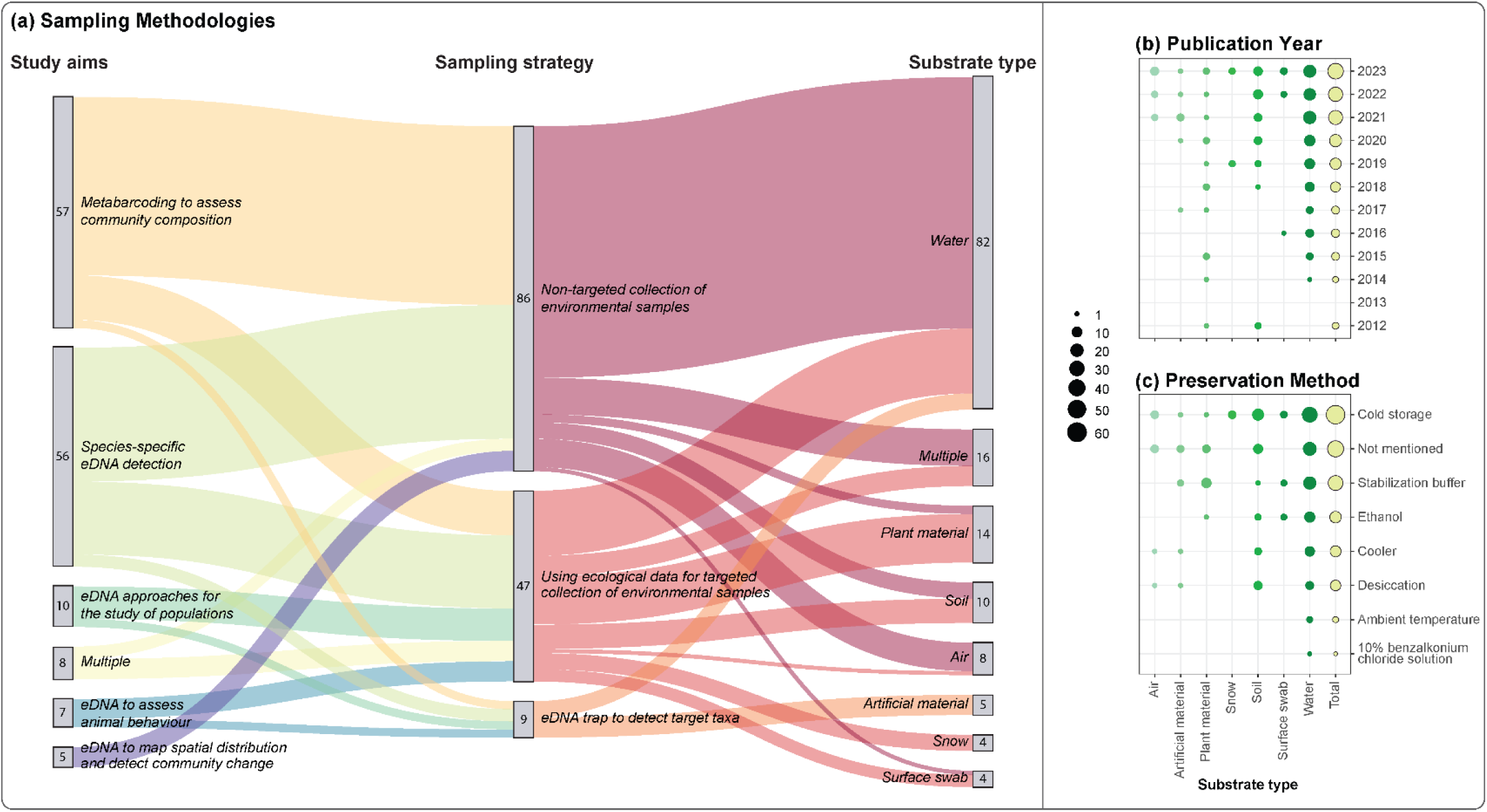
(a) An alluvial plot summarising the study aims, sampling strategies and substrate types used within terrestrial eDNA studies found from the literature search targeting terrestrial vertebrate taxa (N = 143; excluding Lissamphibia) across 2012 – 2023. (b-c) Bubble plots showing the proportion of studies across the substrate types used within terrestrial vertebrate eDNA studies (excluding Lissamphibia) per publication year (b) and preservation method used (c).

#### Sampling strategies, and substrates

To date, numerous sampling strategies and substrates have been utilized for the detection of terrestrial vertebrates using eDNA (Figure 3a). Following our initial literature survey we decided to break down the published sampling strategies into three categories: 1) non-targeted collection of environmental substrate as a non-invasive method to detect target species diversity in an environment (e.g., water collected from rivers for the detection of terrestrial mammals; Broadhurst et al., 2021); 2) specially designed eDNA traps to detect target taxa (e.g., paper-lined nest tubes for the detection of arboreal mammal; Priestley et al., 2021); 3) Using ecological data for targeted collection of environmental samples to detect target taxa (e.g., sampling cover objects for the detection of reptiles or flowers for the detection of nectar feeding species; Kyle et al., 2022; Walker et al., 2022). The non-targeted collection of environmental substrates is currently the most frequently used sampling strategy (86 studies – 60 %) and water the most commonly collected substrate, probably because of the extensive research surrounding eDNA in aquatic systems (Figure 3a; Takahashi et al., 2023; Xing et al., 2022); however, this also reflects the importance on water bodies in terrestrial environments for much of the local biodiversity (e.g., McDonald et al., 2023). Specifically targeted sampling strategies, such as the use of eDNA traps (9 studies) or utilizing ecological data such as behaviour knowledge (e.g., focal water and food sources) and microhabitats, (e.g., tree hollows) to detect target taxa (47 studies) have also been explored. While the majority of research has used either water (94 studies – 66 %), soil (26 studies – 18%), or plant material (14 studies – 10 %), the later part of the decade has seen an increase in studies testing various novel substrates (e.g., air [Clare et al., 2022; Lynggaard et al., 2022] and surfaces [Guthrie et al., 2023; Kyle et al., 2022]) as new environments are explored (Figure 3b). However, only sixteen studies (11 %) have used multiple substrate types, with direct comparisons showing different substrates contain both different levels of eDNA diversity (Allen et al., 2023; Ionescu et al., 2022; Newton et al., 2022; Sales, Kaizer, et al., 2020) and provide varying detection rates (Kyle et al., 2022; Matthias et al., 2021; Sales, McKenzie, et al., 2020).

#### DNA contamination reduction and detection during field sampling

Of the 143 reviewed studies, 74 % (106 studies) specifically addressed decontamination of sampling equipment. Of these, 75 studies stated that sampling devices were “sterile”, “clean”, “DNA free” or “disinfected” without giving further details, two stated that the sampling equipment used was “new” or “freshly opened”. The use of single-use products to reduce contamination was specified 23 times. Where decontamination processes were outlined in greater detail, bleach was the most commonly used approach (42 studies), with UV sterilization (7 studies), flame- sterilization (5 studies), autoclaving (5 studies) and specially designed products such as DNA away® (4 studies) also mentioned. Only one study used alcohol as the sole cleaning agent for sampling equipment, due to problems with bleach supply and transport issues (Lozano Mojica & Caballero, 2021), which is not ideal since alcohol sterilizes but does not necessarily remove DNA contamination (Dickie et al., 2018).

While frequently included throughout the laboratory process, field negative control samples were included in only 61 of the 143 studies reviewed (43 %). Field negative controls can be broken down into two categories: field and equipment negative controls. Field negative controls are collected at each site using exactly the same protocol, equipment, preservation, and processing as field samples, in order to assess whether contamination was introduced during field sampling procedure. Field negative controls were used in 53 studies and all consist of a “target DNA free” sample comparable to the sampled substrate, either collected away from the known location of target species (e.g., Katz et al., 2021), prior to exposure to the target species (e.g., time zero in experimental design; Hunter et al., 2015; Piaggio et al., 2014), or more commonly a known sterile sample (e.g., distilled water) brought to the sample site for the purpose of detecting potential contamination. Alternatively, equipment negative controls, used to assess potential contamination that might be associated with the tools and equipment (e.g., swabbing of muti-use equipment), were collected in only eight studies. Equipment negative controls are vital where multi-use sampling equipment is used (e.g., eDNA traps) to ensure decontamination processes are adequate and no residual DNA is carried over between sampling events.

#### In field sample preservation

The use of cold storage (e.g., ice in cooler, fridge, liquid nitrogen; 47 studies – 32 %) was the most commonly used preservation method, although techniques varied greatly among substrates (Figure 3c). Overall, we found 26 % of studies failed to specify field preservation methods, with no details on time lag between sample collection and cold storage at the laboratory. A further two studies indicated that samples were kept at ambient temperatures while thirteen specified the use of a cooler only. The use of a stabilization buffer (e.g., RNAlater, Longmire’s buffer, buffer ATL; 31 studies – 21%) was also common.

### Five critical considerations for sampling collection

Our review of the literature shows that, while terrestrial vertebrate eDNA biomonitoring has a shorter history than that of aquatic vertebrate species, it has undoubtedly benefited from this previous research testing the limits of eDNA detection and has thus been able to move rapidly into more ecological, hypothesis-driven research involving multispecies metabarcoding. Despite this, there is still a relatively low number of terrestrial vertebrate eDNA studies published relative to aquatic vertebrate eDNA research (74 fish and shark/ray studies published in 2021 alone; Takahashi et al., 2023) when one considers the broad applications of the technique in terrestrial systems. Only the past three years have witnessed a significant increase in relevant publications, possibly suggesting the beginning of an exponential increase. However, with a multitude of study aims, across different environments, and target organisms with different ecologies, the standardization of eDNA sampling approaches for the detection of terrestrial vertebrates is extremely challenging. In the light of this, we have identified five key factors that we believe are essential for future studies (Figure 4). There are, perforce, overlap between some of these factors and, depending on the study, single key factors can be the main aim. However, for most ecological terrestrial vertebrate eDNA studies, all factors are likely to be vital.

**Figure 4.**
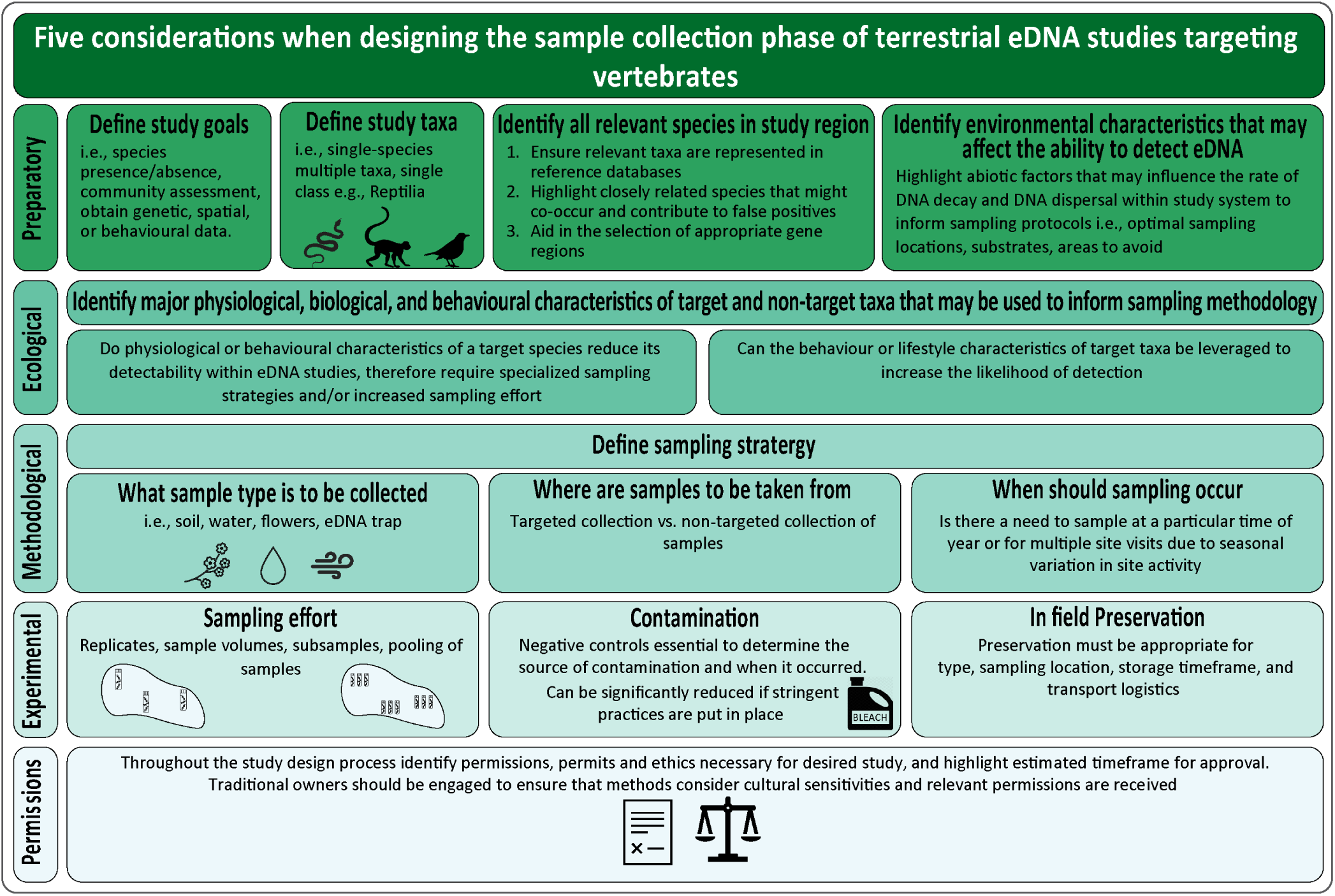
Summary of five considerations that should be addressed when designing the sample collection phase of terrestrial eDNA studies targeting vertebrates.

#### 1. Preparatory Considerations

Before undertaking any terrestrial vertebrate eDNA study, essential preparatory steps will ensure the study’s effectiveness and accuracy. Firstly, defining study goals is paramount i.e., species presence/absence, community assessment, obtain genetic, spatial, or behavioural data. This will ensure appropriate study designs are implemented. For instance, while soil and water have been widely used for community composition assessment (e.g., Leempoel et al., 2020; Mas-Carrió et al., 2021), their suitability for generating population genetic data may be limited (although see J. A. Farrell et al., 2022) due to the complexity of assigning sequences to individuals when only short genetic fragments are obtained (Adams, Knapp, et al., 2019). Clearly defining the study taxon (i.e., single-species, multiple taxa, single class e.g. Reptilia) is also crucial: the benefits of modifying the sampling design to specifically accommodate the target taxa is highlighted throughout the literature and it may be necessary to modify the sampling scheme when specifically targeting rare or cryptic species (Priestley et al., 2021), and/or when assessing behavioural knowledge (Rößler et al., 2020; Stewart et al., 2018). Whereas the use of diverse substrate types is beneficial in broad community assessments (Newton et al., 2022).

Secondly, an understanding of the abiotic factors affecting the ecology of eDNA throughout a sample site can ensure that more appropriate study designs are implemented. It is established that environmental characteristics (i.e., temperature, humidity, UV) of a study site will affect the ability to detect eDNA, due to impacts on factors like DNA degradation. These environmental conditions inform the ‘window’ between DNA shed by an organism and its detection in a laboratory, and thus the limits of applications for biodiversity monitoring. However, within the reviewed literature there are few examples of where abiotic factors are fully taken into account, with the most common considerations being associated with temperature and UV exposure (e.g., Priestley et al., 2021; Ryan et al., 2022). Similarly, few studies have yet quantified the rate of terrestrial vertebrate eDNA degradation (see M. J. Farrell et al., 2022; Guthrie et al., 2024; Kucherenko et al., 2018; Ratsch et al., 2020) and dispersal within terrestrial environments (see Klepke et al., 2022; Lynggaard et al., 2022; Newton et al., 2024); however studies targeting other taxa, and within aquatic, ancient and forensic contexts, indicate that a variety of abiotic factors can influence the rate of DNA decay and DNA dispersal within terrestrial systems (Table 1).

Preliminary studies can be undertaken to determine the temporal ‘window’ for eDNA collection by assessing eDNA accumulation and degradation dynamics of target species and sampling environments (e.g., Kucherenko et al., 2018). In doing so, not only will the temporal ‘window’ be determined, but sampling locations can be selected to increase the probability of successful eDNA detection. For example, by determining the most prominent characteristics likely to cause DNA decay (Table 1), measurements (e.g., temperature or pH) could then be made to identify optimal sampling locations, or areas that should be avoided (e.g., highly sun exposed areas). However, it’s important to acknowledge that it may not always be realistic to conduct such preliminary surveys for every study. In such cases, relying on previous results and common sense should guide the sampling approach to ensure the best possible outcomes.

**Table 1.**
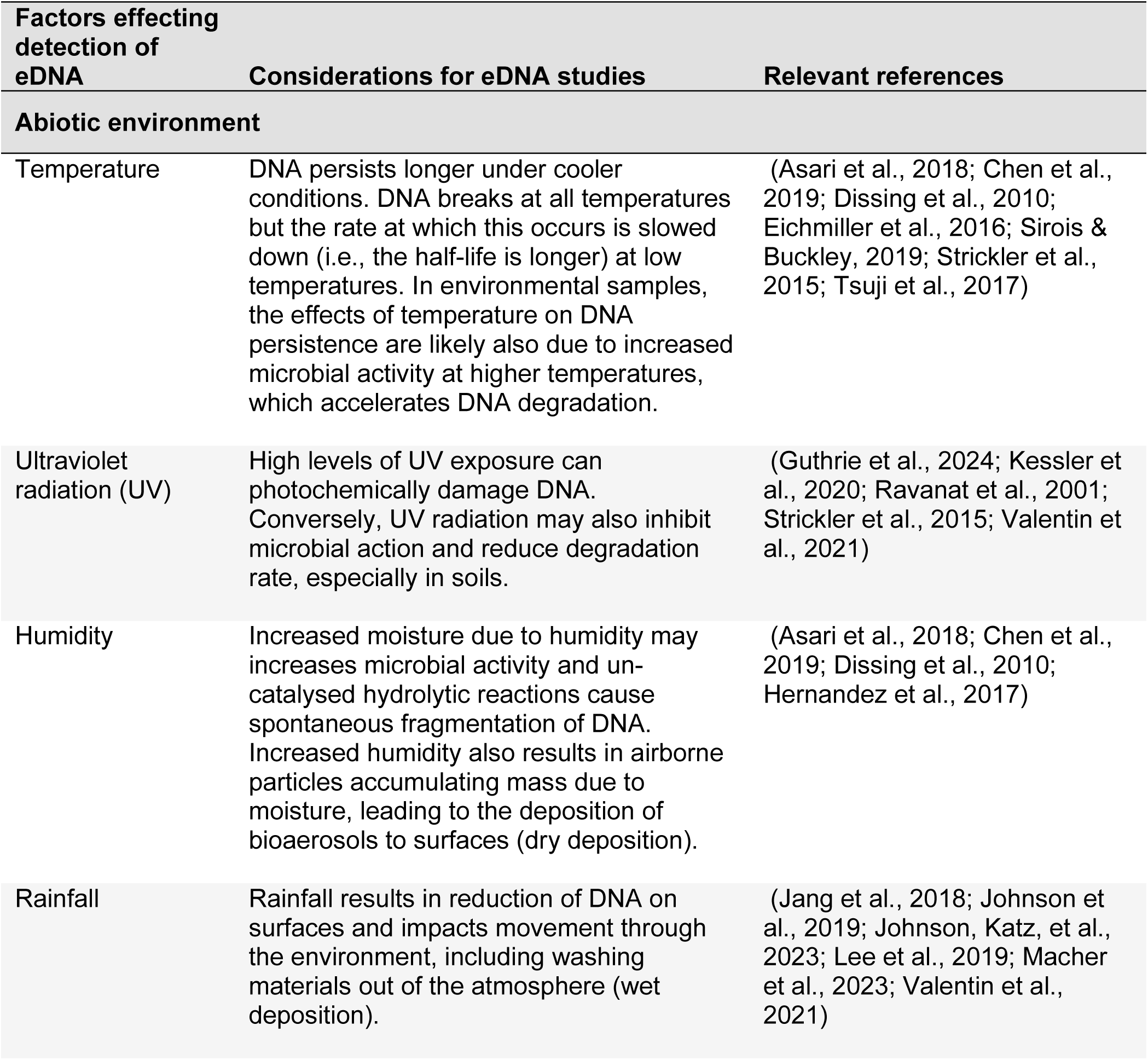
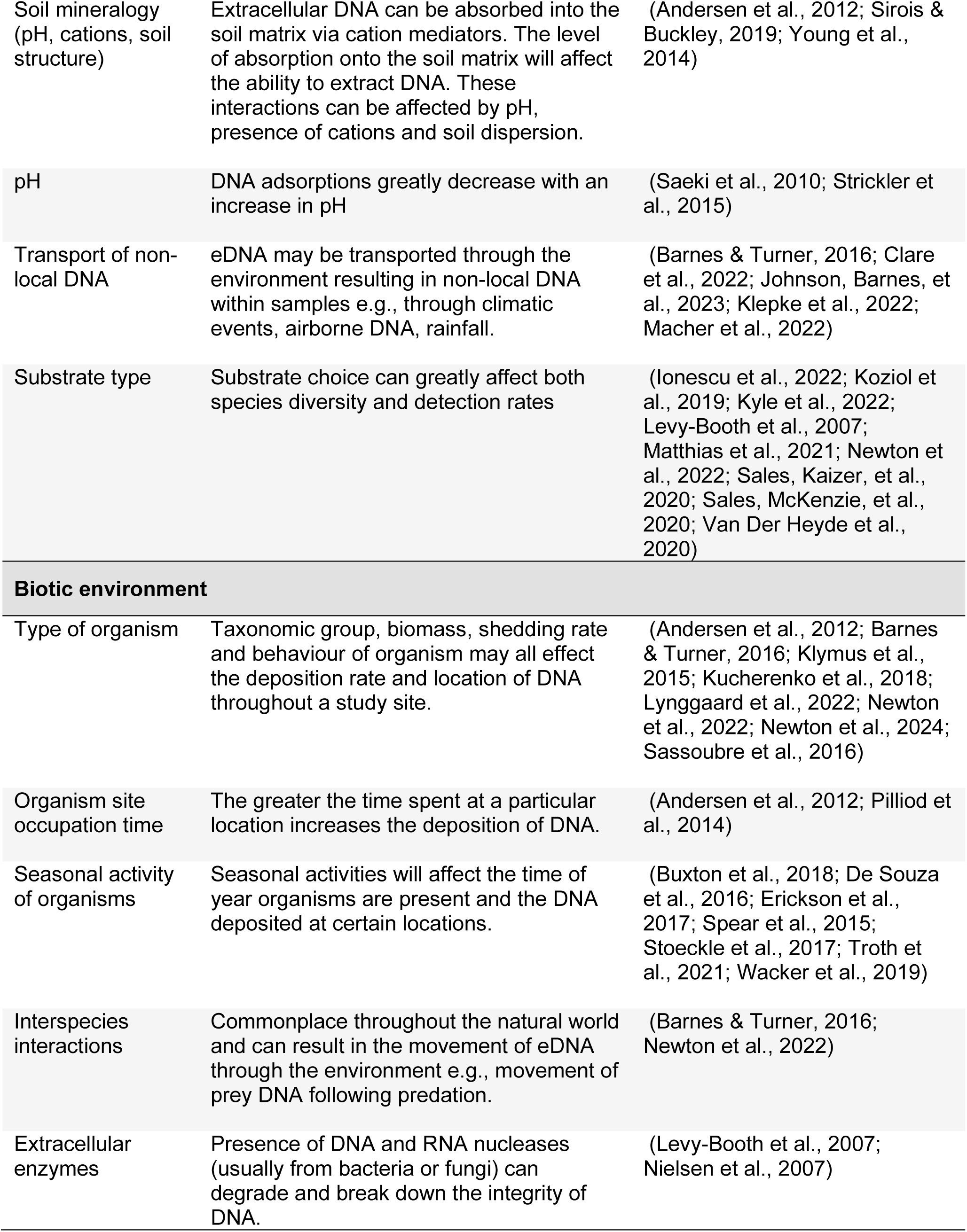
Factors potentially effecting eDNA detection within terrestrial environments and 327 considerations that need to be made when designing terrestrial eDNA studies.

| Factors effecting detection of eDNA | Considerations for eDNA studies | Relevant references |
| --- | --- | --- |
| **Abiotic environment** |  |  |
| Temperature | DNA persists longer under cooler conditions. DNA breaks at all temperatures but the rate at which this occurs is slowed down (i.e., the half-life is longer) at low temperatures. In environmental samples, the effects of temperature on DNA persistence are likely also due to increased microbial activity at higher temperatures, which accelerates DNA degradation. | (Asari et al., 2018; Chen et al., 2019; Dissing et al., 2010; Eichmiller et al., 2016; Sirois & Buckley, 2019; Strickler et al., 2015; Tsuji et al., 2017) |
| Ultraviolet radiation (UV) | High levels of UV exposure can photochemically damage DNA. Conversely, UV radiation may also inhibit microbial action and reduce degradation rate, especially in soils. | (Guthrie et al., 2024; Kessler et al., 2020; Ravanat et al., 2001; Strickler et al., 2015; Valentin et al., 2021) |
| Humidity | Increased moisture due to humidity may increases microbial activity and un-catalysed hydrolytic reactions cause spontaneous fragmentation of DNA. Increased humidity also results in airborne particles accumulating mass due to moisture, leading to the deposition of bioaerosols to surfaces (dry deposition). | (Asari et al., 2018; Chen et al., 2019; Dissing et al., 2010; Hernandez et al., 2017) |
| Rainfall | Rainfall results in reduction of DNA on surfaces and impacts movement through the environment, including washing materials out of the atmosphere (wet deposition). | (Jang et al., 2018; Johnson et al., 2019; Johnson, Katz, et al., 2023; Lee et al., 2019; Macher et al., 2023; Valentin et al., 2021) |
| Soil mineralogy (pH, cations, soil structure) | Extracellular DNA can be absorbed into the soil matrix via cation mediators. The level of absorption onto the soil matrix will affect the ability to extract DNA. These interactions can be affected by pH, presence of cations and soil dispersion. | (Andersen et al., 2012; Sirois & Buckley, 2019; Young et al., 2014) |
| pH | DNA adsorptions greatly decrease with an increase in pH | (Saeki et al., 2010; Strickler et al., 2015) |
| Transport of non-local DNA | eDNA may be transported through the environment resulting in non-local DNA within samples e.g., through climatic events, airborne DNA, rainfall. | (Barnes & Turner, 2016; Clare et al., 2022; Johnson, Barnes, et al., 2023; Klepke et al., 2022; Macher et al., 2022) |
| Substrate type | Substrate choice can greatly affect both species diversity and detection rates | (Ionescu et al., 2022; Koziol et al., 2019; Kyle et al., 2022; Levy-Booth et al., 2007; Matthias et al., 2021; Newton et al., 2022; Sales, Kaizer, et al., 2020; Sales, McKenzie, et al., 2020; Van Der Heyde et al., 2020) |
| **Biotic environment** |  |  |
| Type of organism | Taxonomic group, biomass, shedding rate and behaviour of organism may all effect the deposition rate and location of DNA throughout a study site. | (Andersen et al., 2012; Barnes & Turner, 2016; Klymus et al., 2015; Kucherenko et al., 2018; Lynggaard et al., 2022; Newton et al., 2022; Newton et al., 2024; Sassoubre et al., 2016) |
| Organism site occupation time | The greater the time spent at a particular location increases the deposition of DNA. | (Andersen et al., 2012; Pilliod et al., 2014) |
| Seasonal activity of organisms | Seasonal activities will affect the time of year organisms are present and the DNA deposited at certain locations. | (Buxton et al., 2018; De Souza et al., 2016; Erickson et al., 2017; Spear et al., 2015; Stoeckle et al., 2017; Troth et al., 2021; Wacker et al., 2019) |
| Interspecies interactions | Commonplace throughout the natural world and can result in the movement of eDNA through the environment e.g., movement of prey DNA following predation. | (Barnes & Turner, 2016; Newton et al., 2022) |
| Extracellular enzymes | Presence of DNA and RNA nucleases (usually from bacteria or fungi) can degrade and break down the integrity of DNA. | (Levy-Booth et al., 2007; Nielsen et al., 2007) |

While researchers are investigating the feasibility of targeting longer DNA fragments such as full mitochondrial genomes (see J. A. Farrell et al., 2022), within the reviewed literature, the majority (81 %) of studies targeted a region shorter than 200 bp (Appendix S2, Table S3). Past a certain point, eDNA will become too degraded for a PCR-based approach to work because the strands become too short to bind properly to the primers. By targeting such short regions, the temporal window will be prolonged and increase the possibility of detection. This may allow for sampling in areas exposed to environmental factors such as high temperatures that degrade eDNA, although sampling at multiple time points may be necessary to account for temporally limited DNA signatures (M. J. Farrell et al., 2022). However, re-designing primers for shorter target sequences is not always a viable solution as they must contain enough informative sites for accurate taxonomic identification. For example, Meusnier et al. (2008) showed COI fragments of less than 100bp to clearly reduced the taxonomic resolution. Under circumstances where longer more informative DNA fragments are needed (e.g., eDNA approaches for the study of populations), serious thought into experimental design that considers the places (e.g. microhabitats) where eDNA might be deposited or of specially designed DNA traps (e.g., Emami-Khoyi et al., 2021; Priestley et al., 2021) that may reduce the degradation of DNA will be necessary.

#### 2. Ecological Considerations

The ecological consideration step has two primary goals. First, it aims to elucidate if the study site contains target taxa that are potentially less detectable with eDNA technology, like rare or cryptic species or those with low DNA deposition rates, which would possibly require specialized sampling strategies and/or increased sampling effort (Erickson et al., 2019). Second, it seeks to explore the behaviour and life history traits of target taxa that can be leveraged to increase the likelihood of detection with eDNA.

The ability to successfully recover eDNA deposited by target organisms is governed by two factors: the occurrence of eDNA at the sample site (is the eDNA present at the time of sampling?), and the capture probability (does the sample contain eDNA?; Erickson et al., 2019). However, the exact quantity and quality of deposited eDNA varies greatly across taxa due to physiological, biological, and behavioural differences between species (Table 1). Furthermore, although the overall principles for detecting aquatic and terrestrial vertebrates using eDNA techniques are similar, the sampling strategy is often more challenging in the latter due to more heterogeneous substrates and more spatially restricted dispersal of eDNA in terrestrial systems. Therefore, terrestrial vertebrate DNA is more likely to be recovered from the exact locations where it was deposited (although see Klepke et al., 2022).

To the best of our knowledge, no terrestrial eDNA study has managed to detect all vertebrate taxa known to be present at a given site when employing a metabarcoding approach. Mammalian taxa are often detected in greater numbers than other taxa with the physiological characteristics (e.g., body size, body covering, excretion volume) likely having a significant impact on their DNA deposition rates and subsequently detection rates. The impacts of physiological characteristics on eDNA detection are clearly evident in reptiles, which are commonly undetected in eDNA metabarcoding surveys. It is hypothesised that reduced urine production and a hard or keratinized outer layer to prevent evaporative loss might lead to lower eDNA shedding rates in reptiles compared to other vertebrates (Adams, Hoekstra, et al., 2019; Nordstrom et al., 2022; West et al., 2021) highlighting a need for specialized sampling techniques and/or use of different laboratory protocols to accommodate extremely small amounts of target DNA. Conversely, coverings such as fur, easily shed into the environment, may increase the detection probability of species although this is yet to be fully explored. Recently, both Lynggaard et al. (2022) and Newton et al. (2024) have also shown greater biomass to increase the probability of detection highlighting the importance of considering methods that increase the detection of smaller taxa and/or those with low abundances.

Recent research using ecological data such as knowledge on behaviour patterns and micro habitats to inform targeted sampling (see Newton et al., 2022), and eDNA traps (see Emami-Khoyi et al., 2021; Priestley et al., 2021), suggests that increased understanding of how a species uses a site (e.g., breeding, foraging, seasonal changes, habitat use) can guide sampling strategies (e.g., timing of surveys, substrates, sample locations) and thus improve eDNA detection probabilities. For instance, studies have targeted tree hollows (Newton et al., 2022; Ryan et al., 2022), burrows (Kucherenko et al., 2018), focal water and food sources (Ishige et al., 2017; Mas-Carrió et al., 2021; Newton et al., 2023; Rodgers & Mock, 2015), artificially designed habitats (Kyle et al., 2022; Priestley et al., 2021), and animal tracks (Franklin et al., 2019; Hellström et al., 2023; Von Duyke et al., 2023).

Non-target organisms that are present at the sampling location and their effects on the presence of target eDNA must also be considered. Interspecies interactions such as predation are commonplace throughout the natural world and can result in the movement and introduction of non-local eDNA (Barnes & Turner, 2016). While this cannot be completely avoided the impact may be reduced by sampling in locations that may accumulate non-local eDNA such as dens or nests of predator species. However, if this is unavoidable or these locations are chosen specifically due to their likelihood to contain high quality/quantity of eDNA (e.g., tree hollows used by multiple species; Newton et al., 2022), interpretation of the results must be carefully considered within this biological context.

#### 3. Methodological considerations

When sampling for terrestrial vertebrate DNA throughout the environment the ultimate goal is to identify what substrate is to be collected, where it should be collected from, and when sampling should occur, in order to fulfill the study aims. Within the reviewed literature, sampling strategies can be broadly grouped into two categories: the collection of ‘true’ environmental samples (e.g. soil, plant material, water sources); and targeted systematic sampling, where an eDNA trap (i.e., DNA traces from artificial prey or paper-lined survey tubes to capture a DNA; Priestley et al., 2021; Rößler et al., 2020) is set up in the environment to attract target taxa. While environmental samples have been used more extensively than targeted systems utilizing artificial substrate types (Figure 3a), the choice of which should be driven by the research aims, target taxa and sampling location.

Among the ‘true’ environmental substrates that have previously been used in eDNA studies to target terrestrial vertebrates are soils (e.g., Andersen et al., 2012), focal water sources (McDonald et al., 2023), salt licks (e.g., Ishige et al., 2017), air (e.g., Clare et al., 2022) and plant material (e.g., Nichols et al., 2012). Many ‘true’ environmental samples can be considered indirect sources of eDNA as they rely on an intermediate ‘vector’ (e.g., flower parts) to first collect vertebrate DNA (Garrett et al., 2023; Newton et al., 2023). Consequently, while effective in capturing targeted taxa, these approaches may inherently limit the diversity of species detected. Alternatively, more direct sources of eDNA, such as air, may provide a more unbiased approach (Clare et al., 2022; Garrett et al., 2023; Lynggaard et al., 2022), due to their ability to collect genetic material from organisms that may not interact with traditional vectors. Allowing for the capture of DNA from a broader range of organisms.

Targeted systematic sampling using eDNA traps closely resembles well-established survey techniques such as camera trapping and may allow for more accurate spatial and temporal information. This is due to a higher likelihood of exclusively detecting those species that are currently present and have directly interacted with the eDNA traps during deployment. These traps have been designed to lure target species using baits (Emami-Khoyi et al., 2021), artificial water (Kim et al., 2022; Williams et al., 2018), artificial prey (Rößler et al., 2020), or artificial habitats (e.g., nest tubes/boxes; Priestley et al., 2021). For example, Priestley et al. (2021) successfully detected the rare hazel dormouse (*Muscardinus avellanarius*) by swabbing for DNA deposited on paper-lined nest tubes deployed near permanent nests. While less explored in the current literature target taxa may also be directed/funnelled into the eDNA trap to increase DNA capture (Burns et al., 2020), similar to the widely used pitfall or non-invasive hair traps. While there are a variety of application for eDNA traps, their efficiency heavily depends on their ability to either attract the desired species or to be placed in areas frequented by those species, and effectively capture DNA. Consequently, careful consideration of the behaviour and lifestyle of target taxa (see above) becomes crucial when considering trap design, placement, and deployment, as it will significantly affects the likelihood of successfully eDNA capture.

It is also important to recognize that the choice of any substrate can greatly affect the ability to detect organisms (Koziol et al., 2019; Ryan et al., 2022; Van Der Heyde et al., 2020). Although the majority of studies (∼ 90%) have relied on a single substrate, research that has compared multiple substrate types shows varying levels of eDNA diversity across substrates, (Allen et al., 2023; Ionescu et al., 2022; Newton et al., 2022; Sales, Kaizer, et al., 2020) as well as varying detection rates (Kyle et al., 2022; Matthias et al., 2021; Sales, McKenzie, et al., 2020). For example, Kyle et al. (2022) found that for the detection of reptiles’ swabs of cover objects had a higher detection probability than that of soil samples. Also, the characteristics of a substrate such as proportion of organic matter, pH levels and clay content in soil can influence inhibition in the extraction and/or PCR reaction (Sagova-Mareckova et al., 2008; Young et al., 2014) and may therefore play a role in varying detection rates. Specific substrate may only be available in certain microhabitats (e.g., water bodies) at a study site, or only available at certain time of the year (e.g., seasonal flowering plants). Ultimately, while many potential sources of eDNA are available, pilot studies are essential for each new substrate type to determine its ability to detect the desired diversity in different ecosystems. Finally, given the potential for both eDNA and conventional methods to have biases towards particular species, (Leempoel et al., 2020; Newton et al., 2023; Ryan et al., 2022) how eDNA can complement, rather than replace, conventional methods with complementary strengths should be considered to increase monitoring success.

When selecting an appropriate sampling strategy, it is also essential to consider the environmental conditions specific to each new terrestrial environment that may influence factors such as DNA degradation rates (Table 1). For example, soil has successfully been used to detect vertebrates in colder environments where DNA persists long enough for detection (Andersen et al., 2012; Kjær et al., 2022). However, Van Der Heyde et al. (2020) found that soil performed poorly in hot Mediterranean and hot desert climates with no soil samples tested successfully PCR-amplifying vertebrate eDNA. Consequently, in regions with high temperatures, soil samples may have limited use for vertebrate surveys – at least when based on traditional PCR methods. Depending on these environmental characteristics and the target species, it may be beneficial to sample more stable environments like log hollows (e.g., Ryan et al., 2022) or utilize eDNA traps containing protect from the elements.

At the outset, it is imperative to define the sampling universe, which represents the area from which samples are intended to be representative of. Furthermore, where samples are to be collected from within this universe must also be determined. The choice of sampling location can be chosen using a targeted or non-targeted approach. Targeted approaches may utilize factors such as behaviour patterns, micro habitats or eDNA traps. For example, Ishige et al. (2017) targeted rich natural saltlick sites used by animals to supplement their nutrition, for the detection of local mammal species. This approach is restricted only to those species that interact with those environments or substrates, as such are biased in the species detected. However, this may be beneficial and can provide specific data to answer ecological questions (e.g., classification of tree hollow users; see Newton et al., 2022). Additionally, targeting areas with potential elevated DNA concentrations, for example areas where DNA accumulation may be increased or DNA degradation reduced, such as log hollows used by vertebrate species (see Ryan et al., 2022), can also increase the possibility of detecting target taxa. These techniques may be most effective for the detection of cryptic or rare species, and obtaining population-level genetic information from eDNA samples where quantity or quality of recovered DNA is essential.

Non-targeted sampling, achieved through either objective means (where specific locations are explicitly specified) or subjective methods (where general guidelines are provided, and the exact locations are chosen by the researcher) can provide more reliable and valid data where a broader environmental assessment is the goal. Under this scenario widely distributed environmental substrates such as water, air and soil may be more favourable, all of which have been used to assess community composition involving multiple species (Figure 3a). The choice of these sampling points will dictate how scientifically sound the research will be and, therefore the ability for the research to be replicated and broader conclusions drawn. These critical decisions are thoroughly explored in a recent review (Dickie et al., 2018). Alternatively, while yet to be fully explored, a combination of non-target sample collection for broad species assessment and targeted approaches to detect cryptic and species with low DNA deposition rates may provide more comprehensive datasets.

Due to seasonal variation in site activity (such as migration and breeding activity) there may also be a need to sample at a particular time of year or to conduct multiple site visits (De Souza et al., 2016; MacKenzie & Royle, 2005). For example, De Souza et al. (2016) found that eDNA detection probability for the flattened musk turtle (*Sternotherus depressus*) to be strongly affected by seasonal sampling, with eDNA detection probability highest in the cool season. These results are consistent with the known life history of the species and highlight the value of incorporating biological knowledge into the timing of eDNA sampling.

#### 4. Experimental considerations

After selection of an appropriate sampling strategy and substrate, ensuring an adequate level of field replication becomes essential to enhance confidence in research outcomes. Equally important is vigilance in prevention and detection of DNA contamination, which demands a rigorous approach to maintaining the integrity of collected samples. Furthermore, the preservation of these samples stands as a pivotal consideration, as their condition over time impacts the accuracy of subsequent analyses.

The number of field replicates needs to be established. While greater field replication will often increase the likelihood of detecting the target taxa (Furlan et al., 2016; Willoughby et al., 2016), over-replication is costly. Our review of the literature indicates that a ‘one size fits all’ approach to terrestrial monitoring using eDNA is not possible. The optimal level of sample replication will be dependent on the substrate type, target taxa, and the environment context, all of which, as discussed above, influence the quantity of eDNA present both at the study site (detection probability) and within samples (capture probability). For instance, when detecting species at the edge of their range, rare species (e.g., detection of endangered New Mexico meadow jumping mouse Zapus hudsonius luteus; Priestley et al., 2021), or species with low DNA deposition rates (e.g., reptiles) then more replicates or larger sample sizes are needed. The necessary replication levels are also influenced by the study’s objectives and size of sample universe. Large scale biodiversity assessments often require more extensive survey efforts compared to targeted sampling approaches. Recent research has also highlighted the need to consider temporal replication, especially when sampling in areas that might have a rapid DNA degradation rate (M. J. Farrell et al., 2022).

To mitigate against false negatives, collecting multiple samples per site, and potentially multiple subsamples (non-independent samples taken from sample area and ‘pooled’ to form a sample), may be necessary to minimise the impacts of biological variance found in terrestrial systems (Dickie et al., 2018; Erickson et al., 2019; Ficetola et al., 2015). While there are many possible choices, with all decisions trade-offs will be seen. For example, while pooling samples can increase the scale of analysis, it may reduce the ability to precisely link species to specific locations. Ultimately, while reducing the number of samples or subsamples within sample sites might lower the cost and complexity of a study, it could diminish the capability to capture all biodiversity present, identify biologically meaningful differences, or measure spatial variability at a finer scale (Deiner et al., 2015; Dickie et al., 2018; Hermans et al., 2018).

To estimate the number of samples that are required per site to capture the DNA of target taxa, researchers can use predictive models such as *a priori* occurrence models when the detection probability of target species is known (Erickson et al., 2019), or asymptotic richness estimators (Gotelli & Colwell, 2001). For example, Erickson et al. (2019) utilized data from previous aquatic eDNA studies to show that when detecting common aquatic species, ≤15 samples per site is acceptable. However, when looking for rare species and additional 30 to 75 samples per site are required. They also found that in most cases, ∼30 water samples per site was needed to detect their target taxa more than 95 % of the time. Further research is needed within terrestrial systems and substrates to determine optimal replication level in terrestrial systems. If the detection probability of species cannot be obtained or reasonably assumed from previous studies, a pilot study used to conduct a power analysis would be beneficial. Alternatively, an iterative approach could be used whereby following sampling, outcomes are used to update designs (Goldberg et al., 2016); this would ensure that the experimental designs are appropriate to answer the specific questions asked and detect the desired diversity.

As a corollary to these points, whether the research is exploratory or experimental can also determine many of the sampling parameters. For example, when the objective is to detect a broad array of terrestrial vertebrate taxa using eDNA, a comprehensive approach involving extensive sampling, utilizing multiple substrates, at multiple locations and timepoints might be necessary. Under this scenario, conducting a pilot study for power analysis is beneficial to confirm the appropriateness of sampling methodologies and replication. However, if the aim is to test/trial eDNA methodologies i.e., determine if eDNA methods can detect particular terrestrial vertebrate species using a novel substrate, then fewer samples from one sampling location or samples obtained from a controlled experimental setup may be sufficient (e.g., see airDNA proof of concept study using only 12 sample; Clare et al., 2021).

In the context of terrestrial vertebrate eDNA studies, contamination refers to the inadvertent introduction of foreign DNA into a sample, that is not representative of the local target taxa or environment being studied. It can occur at various stages of the eDNA research process and can have significant implications for the accuracy and reliability of study results. However, limited research has examined background DNA input in the field (see Klepke et al., 2022), despite the increased use of airborne DNA highlighting the undeniable potential of eDNA transport (Clare et al., 2022; Johnson, Barnes, et al., 2023; Newton et al., 2024). Field contamination may occur both across samples (contaminants transferred between sites or samples) and from

The number of negative controls required, and the timing of their collection must be carefully considered. This decision should be guided by the total number of samples and likelihood of contamination, with an increase or decrease in negative controls directly related to one’s ability to determine the source of contamination and when it occurred. For example, Newton et al. (2022) exploring vertebrate use of tree hollow, swabbed multi-use sampling equipment after each field decontamination process. In doing so, they were able to identify contamination on field equipment. However, their stringent field equipment negative controls allowed them to omit only a single species from one sample point due to the detected contamination. Ultimately, finding the right balance between the number of negative controls employed and the breadth of contamination assessment hinges on the specific research context, the intricacies of the study, and the overarching objective of maintaining the integrity of collected data.

Once collected, DNA will continue to degrade – possibly faster than within the in-situ if now exposed to higher temperatures or a less physically/chemically stable environment. This may lead to potential changes in DNA composition and hence a biased reflection of the local terrestrial vertebrate biodiversity. Ideally, DNA extraction should be performed immediately after sample collection to minimize these impacts. However, while mobile laboratories can be utilized, immediate extraction is not always possible (Taberlet et al., 2018). For this reason, preservation techniques are commonly employed. Of the reviewed literature, the use of cold storage (e.g., ice, portable fridge/freezer, liquid nitrogen) and a stabilization buffer (e.g., RNAlater, Longmire’s buffer, buffer ATL) were the most common methods (Figure 3c). However, careful selection of methods is recommended accounting for substrate type, location, transport logistics, and storage timeframe. For example, cold storage can be difficult to achieve in remote field environments where terrestrial vertebrate monitoring is often required. Where it is not achievable, consideration should be made towards the effect this time lag may have on samples, if storage at ambient temperature is acceptable, or a more appropriate preservation approach is available. Techniques such as drying samples using silica gels or preservation buffers such as “Longmire” buffer (Longmire et al., 1997; Taberlet et al., 2018), could present as a more practical alternatives. These techniques enable samples to be stored at ambient temperature, making them particularly useful for remote locations. These methods can also offer advantages when samples need to be transported across significant distances. Notably, the utilization of silica gel in airtight containers as it eliminates the need for flammable or hazardous preservatives, enhancing the safety and ease of transportation (Guerrieri et al., 2021; Taberlet et al., 2018). Furthermore, these methods prove invaluable when researchers are confronted with the challenge of managing large quantities of samples or substantial sample volumes where cold storage is impractical.

When opting for a preservation technique, it is imperative to acknowledge the potential variations in eDNA yield due to storage and preservation. For example, Guerrieri et al. (2021) on the preservation of soil samples suggests that when eDNA extraction is conducted within a few hours of sampling, room temperature storage is sufficient. However, if samples are to be stored for a few days before processing, soil can be stored between 0- 4°C with no significant loss in community composition. However, moderate effects to molecular operational taxonomic unit richness may occur, primarily impacting the detection of rare species. Alternatively, in situations where cold storage is not achievable, drying of soil samples using silica gels maintains ecological signals for at least 21 days, albeit at the cost of detecting rare taxa. Some preservative solutions may also be problematic when used with specific terrestrial substrates. For example, ethanol and RNAlater® have been shown to be ineffective preservatives for environmental samples containing humic acid, potentially due to precipitation and the fixation of nucleic acids to organic compounds causing a reduction in DNA yield during the extraction process (Rissanen et al., 2010). While studies have compared different preservation methods on commonly used eDNA substrates such as water (see Sales et al., 2019) and soil (see Guerrieri et al., 2021), a comprehensive study encompassing a range of potential substrates in various ecosystems is required to shed more light in this area.

#### 5. Permissions and Ethical Considerations

Across the literature reviewed, information on permission for sampling (e.g., research approval, land access, export permits) was common (60 studies); ethics approval was only gained for processes such as the collection of blood and tissue samples conducted in parallel with eDNA studies (see Day et al., 2019; J. A. Farrell et al., 2022; Nichols et al., 2012). When compared to traditional approaches to biodiversity assessment, eDNA surveys sampling abiotic material like soil or water are considered less invasive and non-destructive (to large organisms), and therefore simpler in terms of ethics as they often involve traces of organisms in the environment, rather than handling or trapping of organisms (Fraser & MacRae, 2011). However, ethics approval may still be required where threatened species are targeted or eDNA sampling is to occur on threatened ecological communities. Although, this will vary greatly based on the requirements of local authorities, governments, and research institutions. Where samples are to be transported between countries, permits may be required, potentially unavailable, and affect the feasibility of various substrates. For example, Weiskopf et al. (2018) indicated a three-month waiting period for permits to transport leeches from Bangladesh, used for invertebrate-derived DNA (iDNA) research, and an inability to obtain export permits from both Sumatra and Indonesia. The amount of effort required to obtain permits and ethics approvals is not trivial and must be considered at the beginning and throughout the development phase of all eDNA studies. It would be useful to all researchers in the field if ethics and permit requirements, or lack thereof, were reported in all publications.

In certain situations, broader ethical consideration may also be important, particularly for studies conducted on Indigenous lands or involving culturally important locations and species (Handsley-Davis et al., 2021). For example, in Aotearoa New Zealand where indigenous Māori groups maintain kaitiakitanga (stewardship) over data or resources arising from taonga (culturally important/valued) species. Therefore, this genetic information is considered sacred and subject to protocols derived from tikanga Māori (Māori ethical frameworks) to ensure applications are appropriate. Although eDNA is generally degraded and primarily useful in identifying the presence of an organism at a location, it can reveal information about lands, species and people who occupy or occupied it, which may have inadvertent consequences (Handsley-Davis et al., 2021). For example, eDNA may be used as further evidence to preserve and protect culturally important Birthing Trees used by Aboriginal Peoples in Australia; conversely, the absence of DNA, through degradation or other means, may be used to argue against land rights claims (Handsley-Davis et al., 2021). Under these circumstances, effective communication is required to outline the potential and limitations of eDNA to manage the risk and obtain informed consent from the groups involved. Under all circumstances, the purpose of sample collection and the taxa that may be detected, including both target and non-target, should be outlined to ensure transparency and ethical compliance. This is especially true when using broader sequencing methods such as shotgun sequencing or whole genome sequencing that can reveal all the DNA in a sample, not just the regions needed for taxonomic identification of target taxa (Ng & Kirkness, 2010). Additionally, if further sequencing of stored samples is to be performed, for example applying shotgun sequencing to soil samples initially collected for a vertebrate survey, additional consultation may be required. These deep-sequencing approaches applied to eDNA samples have been shown to readily capture human genomic information termed ‘human genetic bycatch’ (see Whitmore et al., 2023). While human data is rarely the intended target, the potential to inadvertently collect data able to identify and phenotype human individuals may result in future studies utilizing such methods requiring revision by human-focused ethics committees.

### Challenges and Future directions

Many challenges remain with eDNA sampling methods for studies of terrestrial vertebrate species. Interpreting absence remains an enduring issue in eDNA studies and can be influenced by variations in sampling strategy and target taxa, such as the number and location of samples, environmental conditions, biomass, and DNA shedding rates, all of which can influence detection probabilities; however, this challenge extends beyond the realm of eDNA research, as all monitoring data should be interpreted with caution. Future technological advancements will eliminate or reduce many technical sampling issues. For example, purpose-built, portable sampling equipment (e.g., Smith-Root eDNA Sampler) can be designed to optimize sampling speed, replicability and reduce contamination (e.g., Pope et al., 2020; Thomas et al., 2018). Further innovations including 3D printed samplers (Verdier et al., 2022) and autonomous sampling (Truelove et al., 2022) that are currently used in aquatic settings are now being tested in terrestrial studies using air as a DNA substrate (Garrett et al., 2023; Lynggaard et al., 2022), improving the accessibility and scale of sample collection. These innovations, when combined with mobile laboratories, PCR thermocyclers (e.g., Franklin® Real-Time PCR Thermocycler), and sequencing technologies (e.g., MinION nanopore sequencer) would enable rapid on-site biodiversity assessments and detections of invasive species, completely removing the need for storage and preservation. For example, the Smith-Root eDNA Sampler with mobile DNA extraction and qPCR thermocycler to complete eDNA sampling and detect aquatic invasive species in approximately 1 hr in the field (Thomas et al., 2018). This ability to utilize portable eDNA detection tools that deliver rapid results could allow for real-time management actions to be implemented in response to the detection of invasive species (Thomas et al., 2019). However, while these methods are able to identify species in the field and may be of great value for studies conducted in remote areas, or where data is needed quickly (e.g., biosecurity), there are current limitations that need to be considered. It is unlikely these innovative approaches will be available for all substrate types especially those where complex tissue lysing, or extraction techniques are needed to isolate genetic material. However, samples such water or air where tissue lysing is not needed or can be carried out via portable techniques such as physical macerating, and simple methods such as spin-column extractions can be utilised are all viable options for portable laboratory technologies.

Our review of the literature shows that there is no “one size fits all” approach to sampling strategy, eDNA substrate targeted, or sample preservation method. This is best explained by the diversity of study systems, questions explored, and the rapid technological advances in an emerging field. Although there is exciting potential in the eDNA based survey and monitoring of terrestrial vertebrate species, it must be remembered that it is the science that drives the research, not the technology. The issue and the data needed to solve it must remain paramount. Perhaps the biggest challenge for eDNA monitoring in the future are the questions of when and how eDNA is best used (or not), and how it integrates with longstanding approaches including visual observation, trapping and audio or acoustic surveys.

