## Appendix S1 for "Targeting terrestrial vertebrates with eDNA: Trends, perspectives, and considerations for sampling"

Supporting Information

### Appendix S1 – literature search

Search terms

Titles, abstracts, and keywords that contained the terms – “Metabarcod*” OR “Environmental DNA” OR eDNA

With taxa specific search terms

Mammal

“vertebrate” OR “tetrapod” OR “mammal” OR “mammalia” OR “insectivora” OR “mole” OR “shrew” OR “hedgehog” OR “anteater” OR “sloth” OR “armadillo” OR “pholidota” OR “pangolin” OR “chiroptera” OR “bat” OR “carnivora” OR “wolf” OR “coyote” OR “racoon” OR “bear” OR “panda” OR “lion” OR “tiger” OR “feline” OR “fox” OR “leopard” OR “civet” OR “hyena” OR “mongoose” OR “skunk” OR “badger” OR “weasel” OR “otter” OR “rodentia” OR “rodent” OR “rat” OR “mouse” OR “squirrel” OR “capybara” OR “beaver” OR “porcupine” OR “lagomorph” OR “rabbit” OR “hare” OR “perissodactyla” OR “horse” OR “zebra” OR “rhinoceros” OR “tapir” OR “ungulate” OR “deer” OR “pig” OR “hippopotamus” OR “antelope” OR “giraffe” OR “camel” OR “llama” OR “alpaca” OR “sheep” OR “goat” OR “primate” OR “monkey” OR “chimpanzee” OR “gorilla” OR “gibbon” OR “lemur” OR “orangutan” OR “baboon” OR “elephant” OR “hyracoidea” OR “dermoptera” OR “aardvark”

Bird

“bird” OR “avian” OR “palaeognathae” OR “ratite” OR “ostrich” OR “emu” OR “cassowary” OR “rhea” OR “tinamous” OR “neognathae” OR “chicken” OR “duck” OR “geese” OR “swan” OR “pigeon” OR “crane” OR “flamingo” OR “grebe” OR “loon” OR “vulture” OR “hawk” OR “falcon” OR “eagle” OR “owl” OR “parrot” OR “passerine” OR “woodpecker” OR “quail” OR “pheasant” OR “grouse” OR “guineafowl” OR “cuckoo” OR “nightjar” OR “sandpiper” OR “gull” OR “tern” OR “penguin” OR “loon” OR “petrel” OR “shearwater” OR “albatross” OR “cormorant” OR “ibis” OR “pelican” OR “heron” OR “shoebill” OR “kingfisher” OR “bee-eater” OR “stork” OR “finch” OR “sparrow” OR “swallow”

Reptile

“reptile” OR “reptilia” OR “snake” OR “lizard” OR “squamata” OR “tortoise” OR “turtle” OR “testudines” OR “crocodile” OR “alligator” OR “crocodilia” OR “tuatara” OR “rhynchocephalia” OR “terrapin” OR “skink” OR “caiman”
